# Effective holistic characterization of small molecule effects using heterogeneous biological networks

**DOI:** 10.1101/2022.03.23.485550

**Authors:** William Mangione, Zackary Falls, Ram Samudrala

## Abstract

The two most common reasons for attrition in therapeutic clinical trials are efficacy and safety. We integrated heterogeneous data to create a human interactome network that was used to comprehensively describe drug behavior in biological systems, with the goal of accurate therapeutic candidate generation. The Computational Analysis of Novel Drug Opportunities (CANDO) platform for shotgun multiscale therapeutic discovery, repurposing, and design was enhanced by integrating drug side effects, protein pathways, protein-protein interactions, protein-disease associations, and the Gene Ontology, complemented with its existing drug/compound, protein, and indication libraries. These integrated networks were reduced to a “multiscale interactomic signature” for each compound that describe its functional behavior as vectors of real values. These signatures are then used for relating compounds to each other with the hypothesis that similar signatures yield similar behavior. Our results indicated that there is significant biological information captured within our networks (particularly via side effects) which enhance the performance of our platform, as evaluated by performing all-against-all leave-one-out drug-indication association benchmarking. Further, drug impacts on pathways derived from computed compound-protein interaction scores served as the features for a random forest machine learning model trained to predict drug-indication associations, with applications to mental disorders and cancer metastasis highlighted. This interactomic pipeline highlights the ability of CANDO to accurately relate drugs in a multitarget and multiscale context, and paves the way for predicting novel putative drug candidates using the information gleaned from indirect data such as side effect profiles and protein pathway information.

## 1 INTRODUCTION

Drug discovery is the practice of identifying chemical entities with activities useful for treating human diseases. Despite substantial advancements in related technologies, the efficiency of novel therapeutic discovery is severely declining: current drug discovery pipelines on average require over a dozen years and more than two billion dollars to bring a drug to market (Mullard, 2014). The two most common reasons for attrition in drug clinical trials are efficacy, in that the drug is not capable of treating the disease, and safety, in which the compound’s adverse effects do not outweigh any purported benefits, which have contributed to 52% and 24% of phase II and phase III clinical trial failures from 2013-2015, respectively (Harrison, 2016). This culminates in a measly 10-14% approval rate for compounds entering phase I clinical trials (Thomas et al., 2016; Hay et al., 2014; Wong et al., 2019). Drugs fail during trials in part due to traditional methods often employing the ‘magic bullet’ approach, where one biological macromolecule (typically a protein) is targeted by a drug with the hypothesis being that desirably modulating its activity will yield therapeutically beneficial outcomes (Eder and Herrling, 2015). This reductionist view that does not adequately consider the functional impacts of multiple targets on entire biological systems (Maggiora, 2011; Zheng et al., 2013). An example using this rational drug design approach is imatinib, which was developed to successfully target a specific mutated protein in chronic myeloid leukemia. Yet even this drug has since been discovered to have many other protein targets, leading to a wider range of therapeutic uses (Gleich et al., 2002; Joensuu, 2002; Droogendijk et al., 2006; Lee and Wang, 2009). On the other hand, adverse events or side effects are unintended effects of drugs and are evidence of the promiscuous nature of small molecules. These off-target interactions are often neglected during initial drug discovery, which partly explains why these events are the second major cause of clinical trial failure.

Many groups have already successfully implemented systems biology approaches to drug discovery that integrate higher scale and more complex data beyond individual interactions between proteins and small molecules (Akil et al., 2018; Tavassoly et al., 2018; Berg, 2014; Fotis et al., 2018; Csermely et al., 2013; Wallach et al., 2015; Li et al., 2020; Yella et al., 2018). Others have employed computational methods to determine associations between proteins, pathways, or structural elements of compounds to adverse drug reactions (ADRs) to better understand the full spectrum of pharmacological effects that drugs exert on biological systems (Iwata et al., 2013; Liu and Altman, 2015; Yamanishi et al., 2012; Xie et al., 2011; Hart et al., 2016).

Here we describe the use of the Computational Analysis of Novel Drug Opportunities (CANDO) platform for both drug indication as well as ADR prediction. CANDO is a shotgun multiscale drug repurposing, discovery, and design platform whose fundamental tenet or paradigm is to assess the biological or therapeutic potential of small molecule chemical compounds based on their interactions to higher scale entities such as proteins, proteomes, and pathways (Minie et al., 2014; Sethi et al., 2015; Chopra and Samudrala, 2016; Falls et al., 2019; Mangione and Samudrala, 2019; Schuler and Samudrala, 2019; Hudson and Samudrala, 2021; Mangione et al., 2020a; Chopra et al., 2016; Mangione et al., 2020b; Overhoff et al., 2021; Schuler et al., 2021; Moukheiber et al., 2022; Mangione et al., 2022; Bruggemann et al., 2022). Our hypotheses are that compound behavior is describable in terms of their interaction signatures, which are real value vectors representing interactions between a given compound and a library of proteins, pathways, cells, etc. and that compounds with similar signatures will have similar effects in biological systems and therefore can be repurposed accordingly. Novel compounds may also be designed to mimic behaviors observed in desired interaction signatures (Overhoff et al., 2021). The current version (v3) of the platform features thousands of both human approved drugs and investigational compounds and the diseases for which they are indicated/associated, as well as tens of thousands of protein structures from multiple organisms, including *Homo sapiens* and SARS-CoV-2. The platform is primarily benchmarked using a protocol that determines how often two drugs associated with the same indication are considered behaviorally similar based on their proteomic signatures; an overall drug repurposing accuracy is assessed after examining all drug pairs for all indications. Aside from varying the composition of proteins in the interaction signatures (Mangione and Samudrala, 2019), other forms of data used to characterize similarity between compounds in CANDO have included chemical fingerprints, or vectors tallying the presence or absence of chemical substructures (Schuler and Samudrala, 2019), and genome-wide gene expression (Subramanian et al., 2017). Given the substantial heterogeneous multiscale data available in biomedical databases on biological systems and the phenotypic consequences of modulating their activity via small molecules, the CANDO paradigm is well-poised to be used to further elucidate these relationships, especially for small molecule therapeutics and their impacts on proteins, pathways, and diseases.

In this study, a biological network was constructed using the human proteome and its known/predicted interactions with a library of 12,951 drugs/compounds. The network was further supplemented with known interactions between the proteins and their functions in various pathways, as well as their associations to human diseases and entities in the Gene Ontology (GO) (Consortium, 2004, 2015, 2019a). A comprehensive library of ADRs were extracted from both drug labels and adverse event reports (Kuhn et al., 2016; Tatonetti et al., 2012) and were used to describe features and behaviors of drugs. Multiscale interactomic signatures were generated using a vector embedding algorithm (Grover and Leskovec, 2016) from various networks. The multiscale signatures increased the ability of CANDO to assess drug similarity in the context of the indications they are associated with relative to just using their linear signatures. This improvement was particularly striking for networks that included known ADR associations, which implied that side effects contain rich information describing the effects of a drug in biological systems. We then used the multiscale/interactomic network to predict ADRs for all drugs/compounds that were absent from the source databases, and an additional increase in drug repurposing accuracy was achieved after integrating these predicted ADR associations. Further, random forest machine learning models were used to elucidate pathways strongly implicated to contribute to mental disorders and metastatic cancers, which recovered many of the most important pathways known to be associated to both of the respective indications. This implies that antineoplastic and psychoactive drugs may be simultaneously impacting multiple pathways to exert their therapeutic mechanisms.

## 2 RESULTS AND DISCUSSION

Figure 1 provides an overview of our study design. Briefly, multiple heterogeneous databases were integrated into a unified network architecture, which was probed to generate multiscale interactomic signatures for drug-indication benchmarking and prediction of ADR associations, as well as gleaning insight into potential pathways responsible for drug therapeutic mechanisms. A detailed description of the pipelines and protocols is provided in the Methods section.

**Figure 1.**
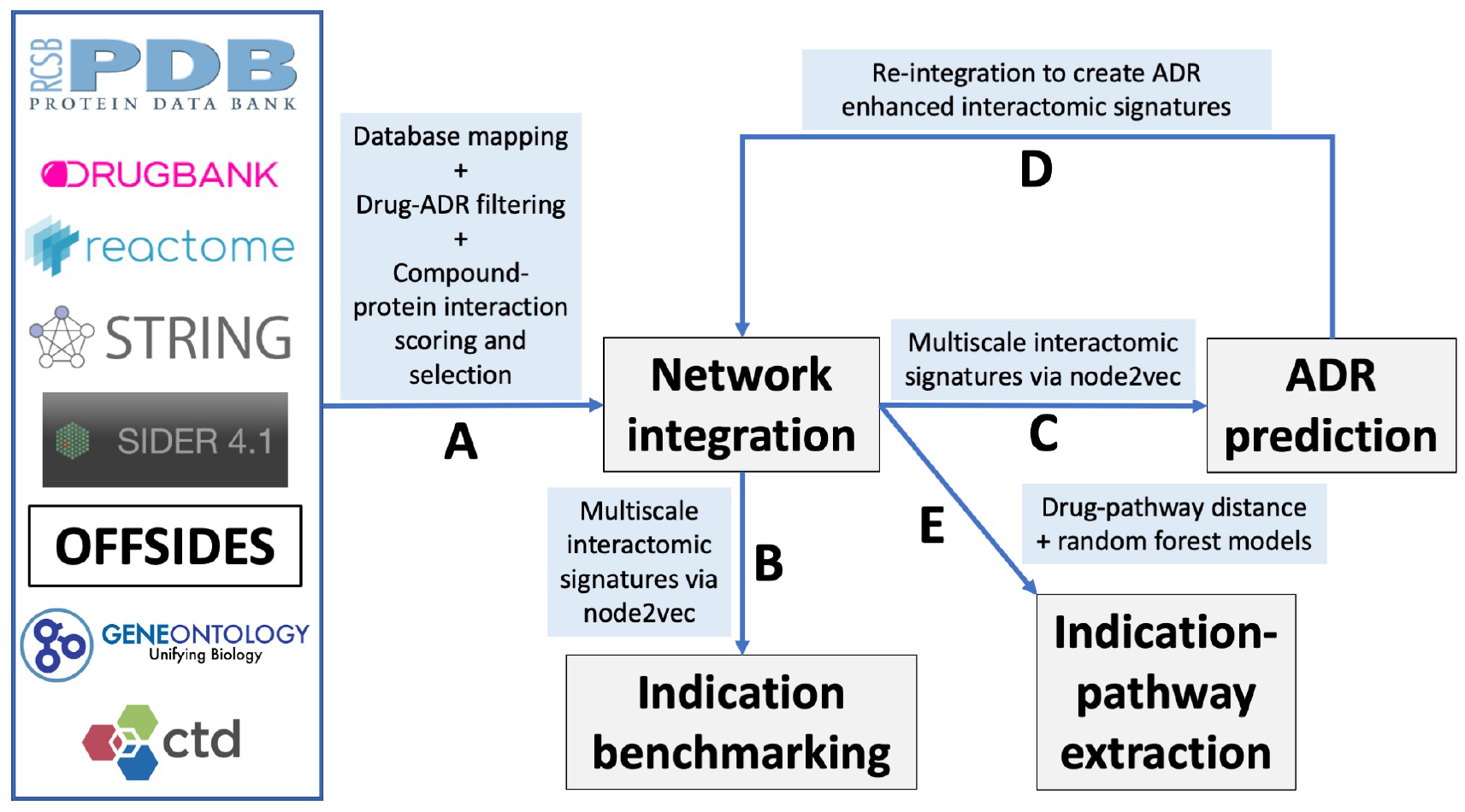
Overview of the CANDO multiscale interactomic signature pipeline and analyses. (A) Multiple biomedical databases were integrated into a unified network architecture after matching the identifiers in these databases to drugs/compounds, proteins, and indications that were already present in the CANDO v2 platform. These databases included DrugBank, the Protein Data Bank (PDB), UniProt, Reactome, STRING, the Gene Ontology, OFFSIDES, and the Comprehensive Toxicogenomics Database. Prior to integration, ADR associations from the OFFSIDES database were filtered to eliminate redundancy with the indication mapping obtained from the Comparative Toxicogenomics Database. Compound-protein interaction scoring were scored using our bioanalytical docking (BANDOCK) protocol. A normalization scheme was also devised to supplement the network with additional interactions (section 4.2). These and other associations between the entities in the heterogeneous databases were used to create an integrated network. (B) A graph feature embedding algorithm, node2vec, was used to create multiscale interactomic signatures from multiple networks. These feature vectors or multiscale signatures were then benchmarked as before (section 4.3) to assess how well they related drugs in the context of the indications for which they are approved. (C) The multiscale interactomic signatures were also used to predict ADR associations for all compounds. (D) These novel ADRs were re-integrated into the network and the resulting multiscale interactomic signatures were benchmarked as in (B). (E) Finally, the distance of a given drug node to each pathway node served as features for input into random forest machine learning models to identify those that are important for predicting drug-indication associations. Coupled with the results of the indication benchmarking, our findings demonstrate that these network-based pipelines are superior to traditional CANDO pipelines for assessing similarity of drug behavior in biological systems.

### 2.1 Benchmarking of networks

All multiscale interactomic signatures outperformed the control proteomic interaction signature matrix in drug repurposing benchmarking accuracy (Figure 2 top). However, the interactomic signatures derived from networks that included ADRs strongly outperformed those from networks excluding ADRs with the mean top10 average indication accuracy of 30.0%.

**Figure 2.**
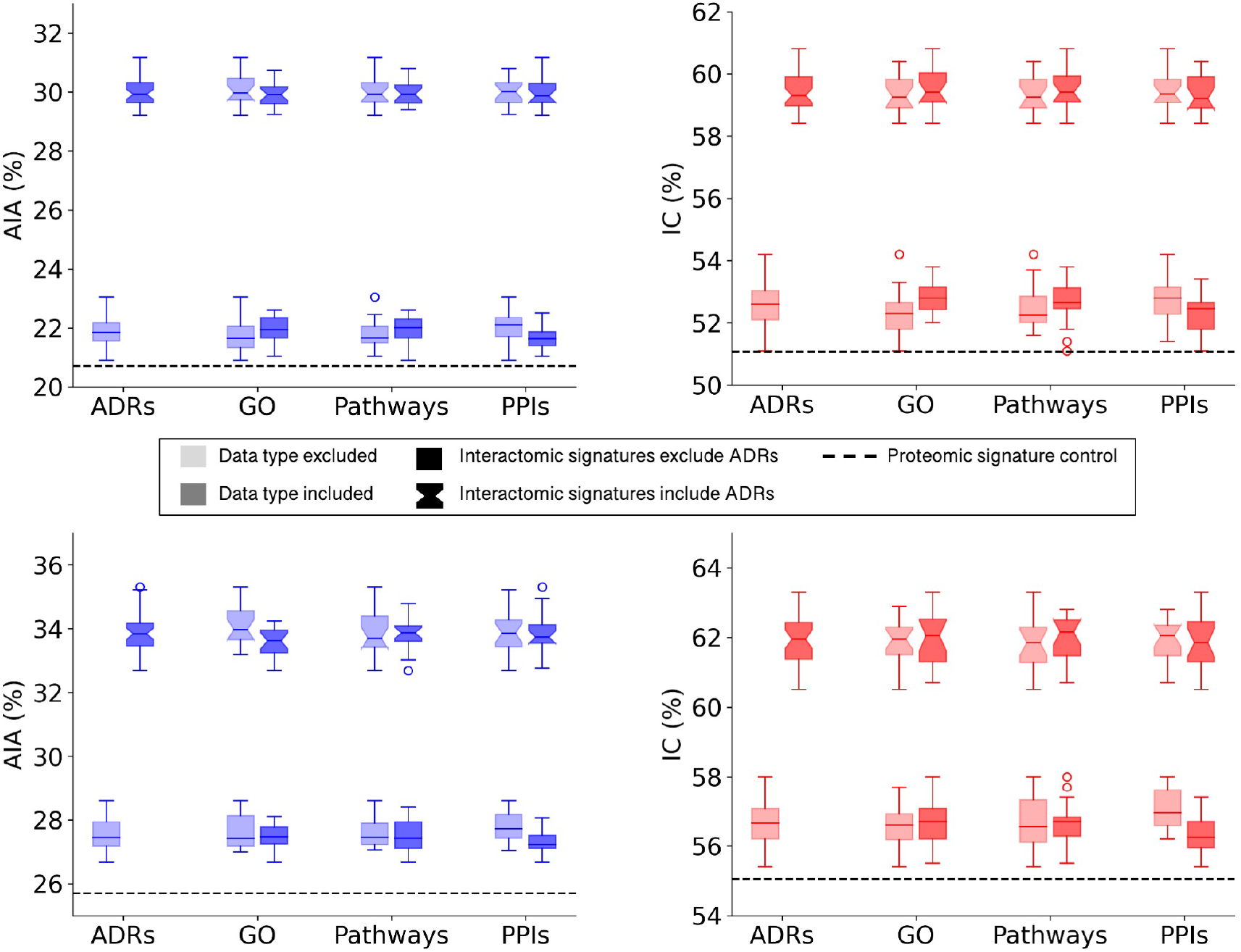
Benchmarking performance of CANDO interactomic signatures with various biomedical data. The difference in benchmarking performance when including or excluding certain biomedical data sources in the multiscale networks, namely adverse drug reactions (ADRs), Gene Ontology (GO) annotations, protein pathways, and protein-protein interactions (PPIs), is depicted for the full drug library (top). Both average indication accuracy (AIA) and indication coverage (IC) followed the same overall trend in which networks that included ADRs (notched boxplots) far exceeded the performance of networks that excluded ADRs (square boxplots). This trend was not observed for the remaining data types in regards to their inclusion (darker) or exclusion (lighter) from the networks. However, all interactomic signatures far exceeded the proteomic signatures control (dashed black line) in performance regardless of the presence of ADRs, indicating the multiscale networks were still accurately capturing drug behavior. The above analysis was repeated with a sublibrary of 1,047 drugs that had at least one ADR associated from SIDER and OFFSIDES (bottom). Similar results were observed with this sublibrary for both average indication accuracy (left) and indication coverage (right), suggesting it is not merely a bias of network architecture. The significant disparity between the performance with and without ADRs implied that ADRs provided rich biological signal for describing drug behavior and motivated us to predict ADRs for all small molecule compounds in the network.

To assess the relative impacts of other relations on benchmarking accuracy that may have been dampened by the significant contribution of the ADR associations, a new compound sublibrary was extracted containing only drugs with at least one ADR and indication association. After benchmarking with only this reduced sublibrary of 1,047 drugs, the interactomic signatures from networks including ADRs still far exceeded the performance of those that did not, though not as dramatically, with a mean top10 average indication accuracy of 33.8% versus 27.6% (Figure 2 bottom). In addition, a slight drop in performance was observed for the signatures that included protein-GO annotations and used DisGeNET thresholds of 0.0 and 0.02 (data not shown), so for subsequent analyses the GO annotations were excluded and the 0.1 threshold was utilized as it provided the greatest coverage of protein associations for indications while still maintaining competitive accuracies. In addition, all network signatures exceeded the performance of the control proteomic interaction signature pipeline, which achieved a top10 average indication accuracy of 25.7% for this sublibrary.

### 2.2 Predicting adverse drug reaction (ADR) associations

The top 100 ADRs were predicted for all compounds using a consensus voting scheme (see section 4.6) and the average *precision@K* at ranks 1 through 100 for all drugs with known ADR associations from OFFSIDES and SIDER was computed (Figure 3). Aside from rank 1, which is the maximum average precision for the top 5% and 10% cutoffs, the peak average *precision@K* was at rank 10 for the top25% cutoff. However, due to a bias discovered in the drug-indication mapping that caused a disproportionate impact on benchmarking accuracy (see below), we selected the top10% cutoff for further analysis due to its better performance at higher ranks.

**Figure 3.**
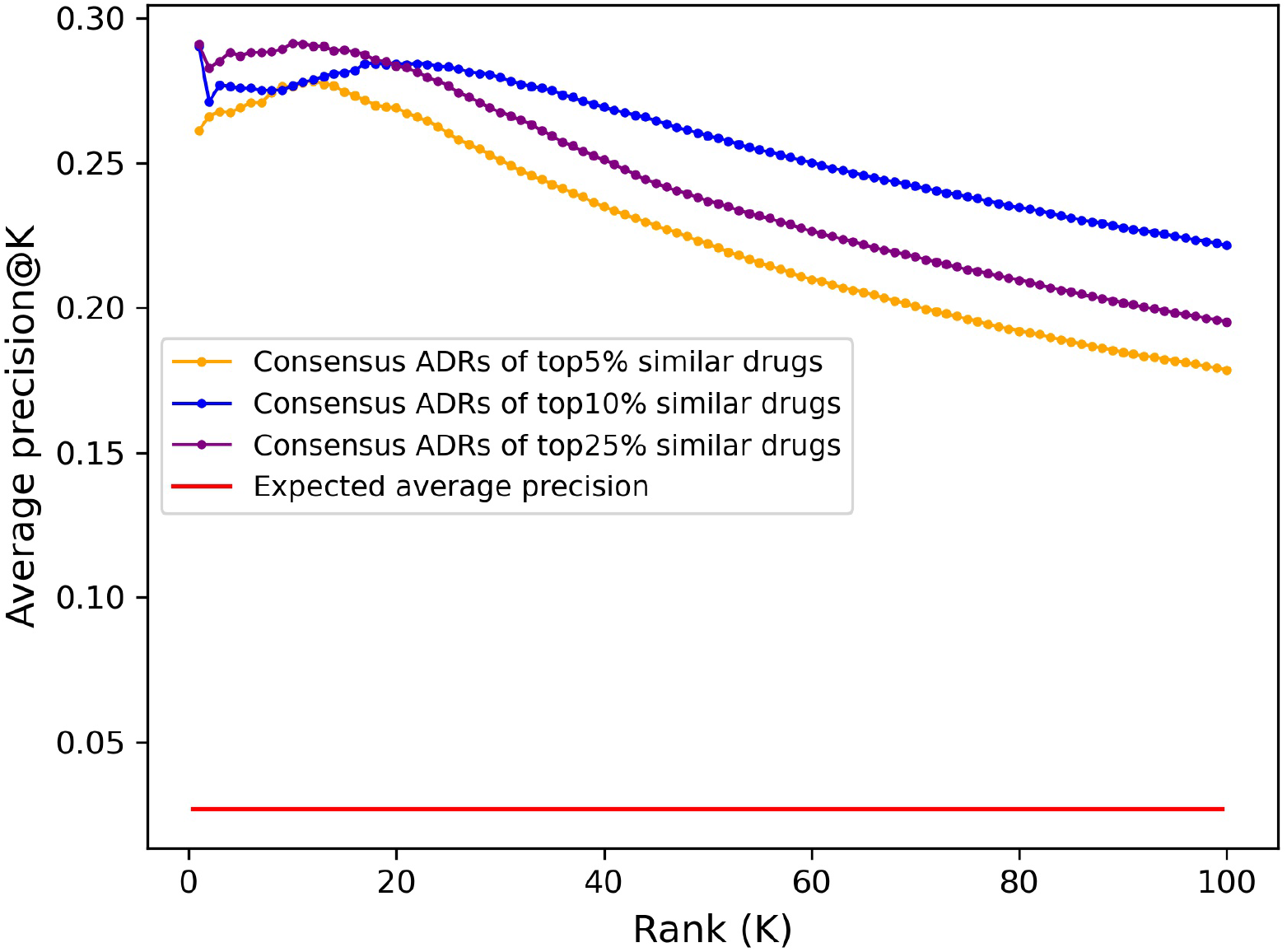
Adverse drug reaction association precision predicted from CANDO interactomic signatures. The top 100 adverse drug reaction predictions for all 12,951 compounds in the CANDO library were generated using a consensus voting scheme based on the hypergeometric distribution. Average precision was calculated at a given rank *K* by averaging the *precision@K* for all drugs with known ADRs (1171). The predictions generated using a cutoff of top 5% (647; orange) had lower average precision across all ranks compared to those generated using a cutoff of top 10% (1295; blue), yet both were 10-fold more precise on average than what is expected at random (red). The predictions using a cutoff of 25% (3238; purple) had higher precision at early ranks, peaking at rank 10, but significantly decreased at progressive ranks. Similarly, aside from rank 1, the average precision of the top 10% lists peaked at rank 18 and steadily declined thereafter, while the top 5% lists peaked earlier at rank 12; this is likely due to the scarcity of drugs with known ADR associations in the full library (9%). The dramatic increase on the expected average precision for all sets of predictions indicated that the interactomic signatures captured compound behavior effectively.

### 2.3 Integrating novel ADR associations

Embeddings were generated from the interactomic networks using the complete compound library, their newly associated top25 or top100 ADRs (from the previous step), protein-protein interactions, and protein-pathway associations, with a DisGeNET score threshold of 0.1. The top25 ADR-enhanced network achieved a top10 average indication accuracy of 28.1%, exceeding that of the top100 ADR-enhanced network of 25.8%; yet both exceeded the aforementioned non-network proteomic signature control performance of 20.7% (Figure 4). The ADR-enhanced networks also outperformed another control matrix that used only the Dice coefficients between drugs in the library as the compound-compound similarity scores, which achieved a top10 average indication accuracy of 23.6% for the approved drug sublibrary. These fingerprint only pipelines typically outperform all other proteomic based pipelines in benchmarking based on previous studies (Schuler and Samudrala, 2019), further indicating the multiscale interactomic signatures were superior at capturing therapeutic signal.

**Figure 4.**
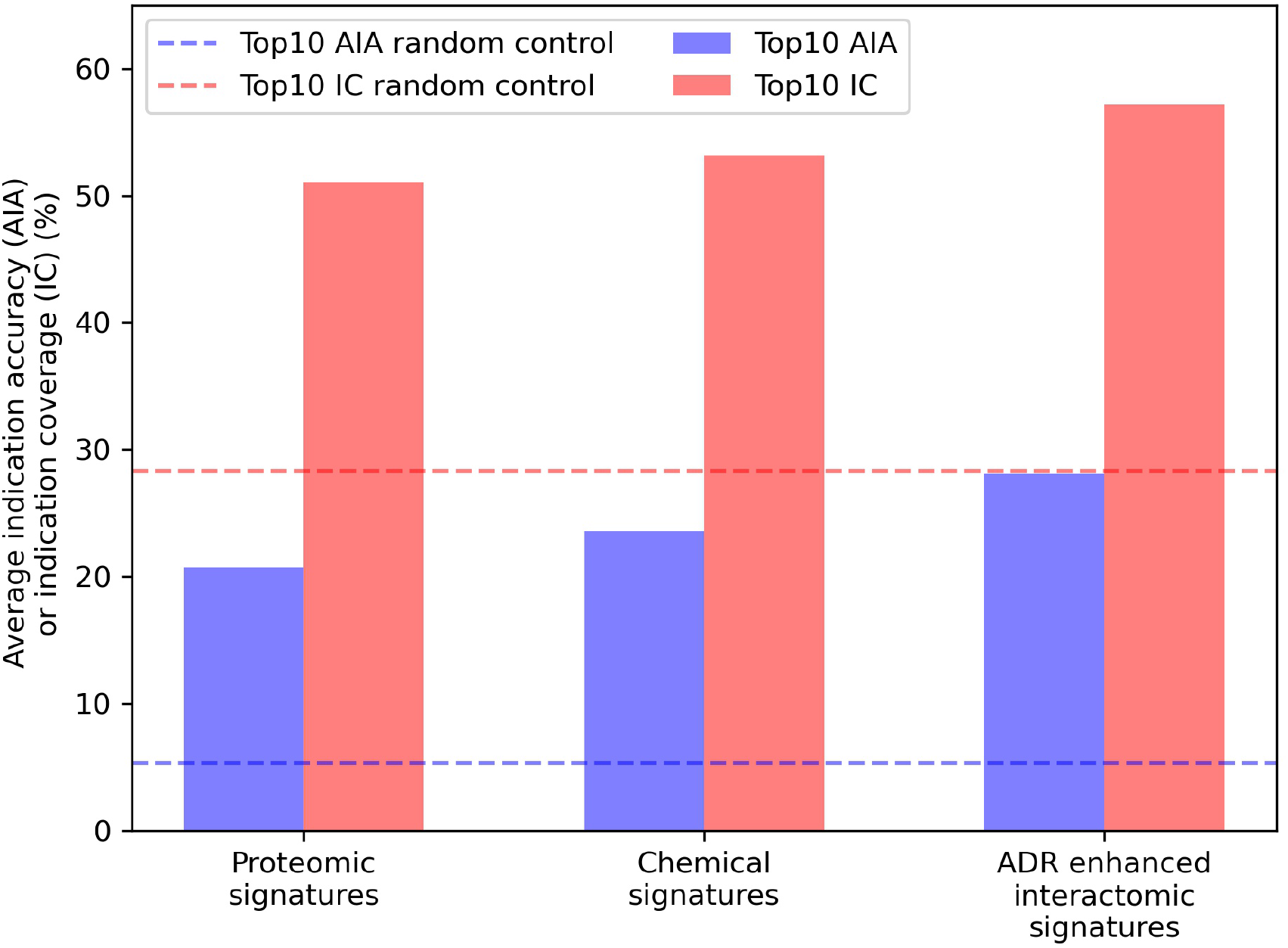
Benchmarking performance of various CANDO pipelines. The ability of different CANDO pipelines to accurately determine similarity of drugs known to treat the same indications was assessed using a hold-one-out benchmarking scheme on a per indication basis. All indication accuracies were averaged to compute the average indication accuracy (AIA; blue), along with counting the number of indications with a non-zero accuracy, or indication coverage (IC; red). The performance of the pipeline using linear proteomic signatures with a library of 8,385 human proteins and the chemical fingerprint pipeline are depicted and compared to the pipeline with human multiscale interactomic networks. The latter was created from continuous-valued low dimensional embeddings based on connectivity of the heterogeneous biological network (see Methods). The multiscale pipeline also comprised a set of the top25 predicted ADR associations for all compounds generated using a consensus voting scheme (see section 4.6). This multiscale ADR association pipeline achieved an average indication accuracy of 28.1% which exceeded the performance of both the proteomic signature and fingerprint pipelines of 20.7% and 23.6%, respectively, and was over five times that of the random control value of 5.3% (dashed blue). This was also true for indication coverage where the ADR-enhanced interactomic signatures achieved 57.2%, amounting to 97 additional indications for which there is a non-zero accuracy (out of 1,588), with an expected random control value of 28.3% (dashed red). This further indicated the multiscale interactomic signatures captured compound behavior more effectively than the less dynamic proteomic and fingerprint signature pipelines. Overall, our results demonstrated that considering interactions between compounds and larger scale entities present in their biological and environmental contexts is necessary for more accurately understanding their therapeutic mechanisms and outcomes.

However, despite featuring ADR associations for all drugs in the CANDO library, the top25 ADR-enhanced interactomic signatures were still outperformed by the signatures generated using the equivalent network with incomplete compound-ADR coverage in benchmarking accuracy (28.1% vs. 30.6%, respectively). This was inconsistent with previous results indicating that ADRs provide rich biological information, so the introduction of the predicted ADR associations should have increased benchmarking performance for the entire approved drug sublibrary. The exact top10 drug-drug-indication hits responsible for the increase in accuracy were inspected; of the hits produced by the incomplete ADR network that were not produced by the complete ADR network, 74.6% were drug-drug pairs where either both *had* or both *did not have* ADR associations (directly from SIDER or OFFSIDES), indicating this pipeline may have preferentially ranked compounds relative to each other based on the sheer quantity of ADRs associated. Given the split in the approved drug sublibrary was essentially even between drugs with known ADRs (1,171) and those without (1,165), this was strong evidence of this network being overtrained.

### 2.4 Analyzing repurposing accuracies for indication classes

In addition to average indication accuracy, both the average pairwise accuracy and indication coverage dramatically increased for the ADR-enhanced networks relative to controls. The average pairwise accuracy reports the percent of successful hits within the top drug-indication pairs relative to all the ones available in the platform (such as top10, top25, etc.). The ADR-enhanced interactomic signatures achieved a top10 average pairwise accuracy of 46.7% compared to the control proteomic and chemical signature (fingerprint) pipeline values of 35.7% and 38.7%, respectively, which constituted an increase of 2,070 and 1,502 successful top10 drug-indication hits, respectively. Accuracies were improved for a total of 616 indications relative to using the proteomic signature pipeline, and for 550 indications relative to using the chemical signature (fingerprint) pipeline (Figure 5). The strongest improvement was in the Neoplasms class (216 indications) in which the multiscale interactomic pipeline successfully improved benchmarking accuracies for 133 (p < 1e-12) and 125 (p = 1e-13) indications compared to the proteomic and chemical signature pipelines, respectively. This strongly indicated the contextual biological information of the network more accurately captured therapeutic behavior compared to the linear proteomic signatures or the protein-agnostic chemical signatures (fingerprints) for complex indications. This was also true for other indications with complex etiologies like cardiovascular conditions and mental disorders.

**Figure 5.**
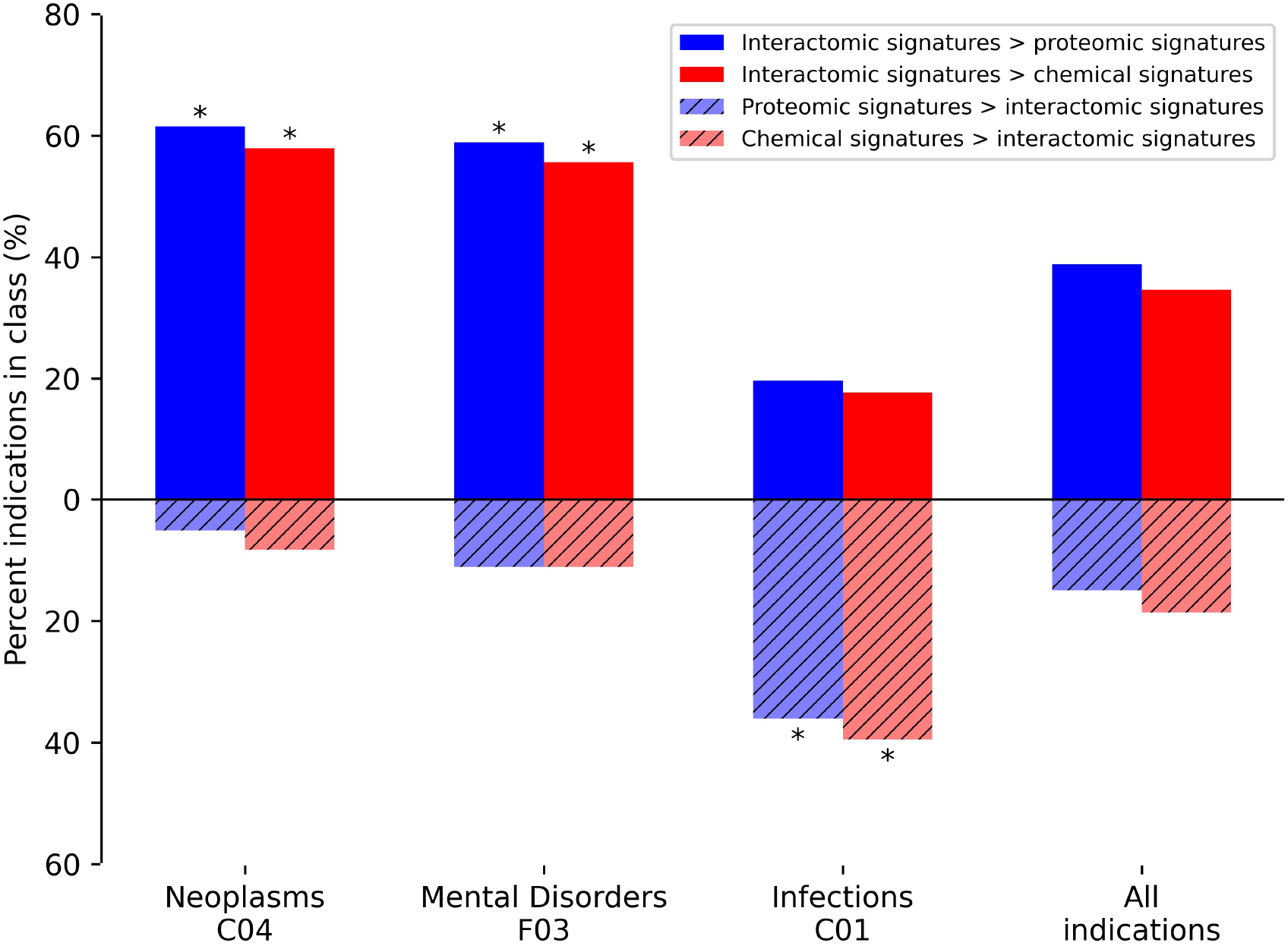
Comparative benchmarking performance by various CANDO pipelines for specific indication classes. The most overrepresented classes of indications, based on their upper level Medical Subject Heading (MeSH) (Lipscomb, 2000)), with changes in benchmarking accuracy for the interactomic signatures compared to both the proteomic and chemical signatures are depicted. The average indication accuracy of the multiscale interactomic pipeline of 28.1% exceeded those of the proteomic and chemical signature pipelines of 20.7% and 23.6%, respectively (section 4.3 and Figure 4). The interactomic signatures definitively outperformed the proteomic signatures (solid blue) and chemical signatures/fingerprints (solid red) for both Neoplasms and Mental Disorders, with significance (asterisks; probability ≤0.005) calculated using the number of indications used for benchmarking (1588), the frequency of each upper level Medical Subject Heading class among those indications, the number of indications belonging to each class with improved scores, and the total number of improved indications using the hypergeometric distribution. However, for the Infections class, the proteomic and chemical signatures (striped blue and red, respectively) performed the best. Given that drugs treating infectious disease typically target pathogenic proteins, which are not included in our human network used to create our interactomic signatures, this result was expected. The proteomic signature pipeline achieving significantly better performance, despite being composed of interactions with human proteins, is a phenomenon observed in previous CANDO publications in which the increase in the total number of proteins in an interaction signature leads to predictable increases in accuracy (Minie et al., 2014; Sethi et al., 2015; Mangione and Samudrala, 2019), most likely due to an increase in the coverage of the protein fold space (Sethi et al., 2015). Overall, the interactomic signatures were able to capture drug behavior in the context of the entire biological system, not limited to a small number of protein targets, as evidenced by the the substantial increases in accuracy for complex diseases like cancer and psychiatric disorders, despite them sometimes possessing diverse and poorly understood etiologies.

Our results provide further evidence that the compound embeddings produced from the networks captured behavioral signal beyond just the scope of the few immediate proteins with which they interact (as in their linear proteomic signature), and were able to capture the context of these interactions relative to the human-based indications with which they are associated. In contrast, the proteomic and chemical signature pipelines achieved higher accuracies for 53 (p = 3e-12) and 58 (p = 1e-10) of the indications in the Infections class, respectively, relative to using the multiscale interactomic signatures (Figure 5). This is likely due to the mechanism of most drugs used to treat infectious diseases (antibiotics, antiparasitics, antivirals, etc) being to directly disrupt pathogenic proteins/processes, none of which are included in the human focused biological networks in this study. Also, the proteomic signature pipeline achieving significantly better performance for Infections indications, despite being comprised solely of human proteins, is a phenomenon observed in previous studies in which the increase in the total number of proteins in an interaction signature, regardless of organismal source, leads to predictable increases in accuracy (Minie et al., 2014; Sethi et al., 2015; Mangione and Samudrala, 2019). This is most likely due to an increase in the coverage of the protein fold space and a consequence of evolutionary conservation between protein structure and function (Sethi et al., 2015).

### 2.5 Associating drugs and pathways

Random forest machine learning models were trained for all indications with at least 50 associated drugs using the distance to a subset of 821 pathways as features for the drugs, and the the resulting vectors were binarized (section 4.7). Among the 60 indications with at least 50 associated drugs, the top five with highest accuracy included “Mental Disorders” (MeSH:D001523), “Migraine Disorders” (MeSH:D008881), “Schizophrenia” (MeSH:D012559), “Dyskinesia” (MeSH:D004409), and “Neoplasm Metastasis” (MeSH:D009362) with average accuracies of 0.83, 0.78, 0.77, 0.77, and 0.76, respectively, over ten iterations of training and testing. The pathway features most important for predicting these drug-indication pairs were extracted for the top two (distinct) indications, “Mental Disorders” and “Neoplasm Metastasis”, with their top five non-redundant pathways listed in Table 1 and Table 2, respectively.

**Table 1.** Pathways most important for predicting Mental Disorders. Pathway importance as extracted from random forest models trained on drugs known to treat Mental Disorders are listed. The models for this indication, which had 54 drugs associated, achieved an average accuracy of 0.83 across 10 iterations of training with a random 90% of the associated drugs as positive samples and testing with the remaining 10% (including an equal number of randomly selected “negative” training/test samples from the rest of the drug library). The ranks of each pathway belonging to each group, their Reactome identifiers, and description of each pathway are listed accompanied by literature references supporting their importance in psychological disorders. These pathways, aside from cytochrome P450 substrate metabolism, are well-known targets of many antipsychotic drugs and provided evidence that the pathways extracted by the random forest models from our interactomic networks were relevant to therapeutic mechanisms.

| Ranks | Reactome identifiers | Pathway group description | # of drugs | Evidence |
| --- | --- | --- | --- | --- |
| 1 | HSA-390651 | Dopamine receptors | 4 | (Ayano, 2016; Ueno, 2003) |
| 2 | HSA-375280 | Amine ligand-binding receptors (GPCRs) | 14 | (Nickols and Conn, 2014; Komatsu, 2015; Komatsu et al., 2019) |
| 3 | HSA-1296071 | Potassium channels | 22 | (Guglielmi et al., 2015; Noh et al., 2019; Judy and Zandi, 2013) |
| 4 | HSA-390666 | Serotonin receptors | 5 | (Ueno, 2003; Iwamoto et al., 2009) |
| 5 | HSA-211958 | Miscellaneous substrates of cytochrome P450s | 54 | (Haduch and Daniel, 2018) |

**Table 2.** Pathways most important for predicting Neoplasm Metastasis. Pathway importance as extracted from random forest models trained on drugs known to treat the Neoplasm Metastasis class are listed. The models for this indication, which had 68 drugs associated, achieved an average accuracy of 0.76 across 10 iterations of training with a random 90% of the associated drugs as positive samples and testing with the remaining 10% (including an equal number of randomly selected “negative” training/test samples from the rest of the drug library). The ranks of each pathway belonging to each group, their Reactome identifiers, and encompassing description of each group are listed accompanied by literature references supporting their importance in cancer. All pathways have multiple studies supporting their relevance, indicating these antineoplastic drugs may be exerting their therapeutic effects via these pathways, serving as a hypothesis generation tool for further validation and also for further multiscale drug development.

| Ranks | Reactome identifiers | Pathway group description | # of drugs | Evidence |
| --- | --- | --- | --- | --- |
| 1 | HSA-156581 | Methylation via S-adenosylmethionine | 63 | (Mahmood et al., 2020, 2018; Parashar et al., 2015) |
| 2; 3; 5; 8 | HSA-9018677; HSA-9018678; HSA-9018682; HSA-9027307 | Biosynthesis of specialized proresolving mediators and maresins | 64 | (Lavy et al., 2021; Vatnick et al., 2016; Serhan et al., 2009) |
| 4 | HSA-5423646 | Aflatoxin activation and detoxification | 62 | (Sudakin, 2003; Marchese et al., 2018; Jackson and Groopman, 1999) |
| 6; 7 | HSA-2142670; HSA-2142691 | Synthesis of epoxy- and dihydroxy-eicosatrienoic acids, leukotrienes, and eoxins | 63 | (Poff and Balazy, 2004; Claesson, 2009; Heyd and Eynard, 2003; Apaya et al., 2020) |
| 8 | HSA-77289 | Mitochondrial fatty acid beta-oxidation | 64 | (Ma et al., 2018, 2021; Liu, 2006) |

## 3 LIMITATIONS AND FUTURE WORK

### 3.1 Sampling of drug-effect associations

Despite the significant improvement in drug repurposing accuracies outlined, the reliance on a library of drugs with known ADR associations amounting to less than one-tenth of the total compounds analyzed in this study may have created a bias for the predictions made for those compounds. In general, a major problem with public data sets in drug discovery is that coverage of the sample space is far from comprehensive. Further, a significant amount of information present from the OFFSIDES project had to be removed due to an inability to confidently determine that the “ADR” associations were not a known symptom/sign of the disease for which the drugs are common treatments. Future work that painstakingly utilizes a biomedical entity hierarchy even more so than was done with SNOMED CT in this study, or assesses the co-linearity of ADRs for drugs indicated for the same indications, will garner much more data crucial for understanding compound behavior.

Related to this, the lack of negative samples for both ADRs and indications likely hinders the performance of the machine learning models used in this study due to an imperfect random selection of compounds for training. Curating a library of negative drug-effect associations may be possible for one or a handful of indications, but producing a library on the scale of the mapping provided by the CTD will require extensive automated parsing of the biomedical literature and clinical trial data, which we will address in a future study.

Further, the tremendous impact the ADRs had on benchmarking indication accuracies, and therefore describing drug behavior in totality, was likely due to the sheer number of compound “effects” that were added by the ADR associations being nearly six times greater than those available in the indication mapping from the Comprehensive Toxigogenomics Database (CTD) as well as an increase in granularity for phenotypic outcomes after exposing these compounds to biological systems. For example, the ADRs can range anywhere from routine diagnostic findings, such as “Blood urea abnormal”, which is likely due to impacts on one or a few protein pathways, to higher level ADRs, such as “stomach cramps” or “headache”, which are likely due to multiple contributing pathways, all of which the interactomic signatures were able to capture. This is highly valuable because an important limitation of most drug-indication prediction methods is the lack of a significant amount of positive samples from which we may glean information about their mechanisms. This will be explored in a future work in which the pathways responsible for causing ADRs will be extracted in a similar manner as they were for the indications in this study (see section 4.7), and clustered based on what pathways are seemingly contributing to those outcomes. We believe the interactomic signatures using these ADRs took advantage of this granularity and learned compound behavior at multiple levels of complexity, showcased by their enhanced ability to relate drugs approved for the same diseases.

### 3.2 Assumptions of network architecture

Based on the binary architecture of the network, it is not obvious which compound-pathway impacts are agonist or antagonist. The same holds true for the docking protocols used for supplementing the network in addition to the known compound-protein interactions (section 4.2). This is already being addressed in a future work in which the LINCS1000 and Connectivity Map gene expression databases are being integrated to better determine whether a compound activates a pathway based on the gene expression of its members, or vice versa (Subramanian et al., 2017; Lamb et al., 2006). The present vector embedding algorithm used here (Grover and Leskovec, 2016) will likely be inefficient for analyzing these relationships, which is why other network based models such as TransE and its derivatives, which specifically handle edge types differently and have been applied to biological data (Choi and Lee, 2021; Peng et al., 2020; Ye et al., 2019; Tu et al., 2017), are being explored.

The diminished ability of the interactomic signatures to capture the therapeutic effects of drugs used to treat infections is a consequence of their protein targets being absent from the network. Future work is planned where various organism-specific biological networks with known pathogen-human protein-protein interactions will be constructed and probed in the same interactomic manner as the human-only network used in this study. The promise of this method extends beyond just direct inhibition of the pathogenic proteins as well; consider a drug such as dexamethasone, which is now a primary treatment for hospitalized COVID-19 patients: the mechanism of this drug is to reduce the severe inflammation cascade (cytokine storm) resulting from viral processes, but is not a direct inhibitor of SARS-CoV-2 proper (Group, 2021). A well-constructed network connecting known viral proteins to inflammatory pathways in humans is made possible by our approach to decipher these relationships and predict effective host-based treatments in addition to direct viral inhibition Draghici et al. (2021); Maxwell et al. (2021).

In the networks used for this study, there is a lack of pharmacokinetics/pharmacodynamics measurements that further describe compound behavior. Despite the interactomic network still producing impressive results without this data, future work will explore dimensions such as drug dose, concentration over time, in different cells and tissues, metabolites, etc. that ideally yield realistic interactions from the analyses.

## 4 METHODS

### 4.1 Heterogeneous biological data collection

Numerous biological databases and tools were utilized to construct the heterogeneous networks used in this study. An overview of the network architecture is depicted in Figure 6 and a detailed description of the data curation follows below.

**Figure 6.**
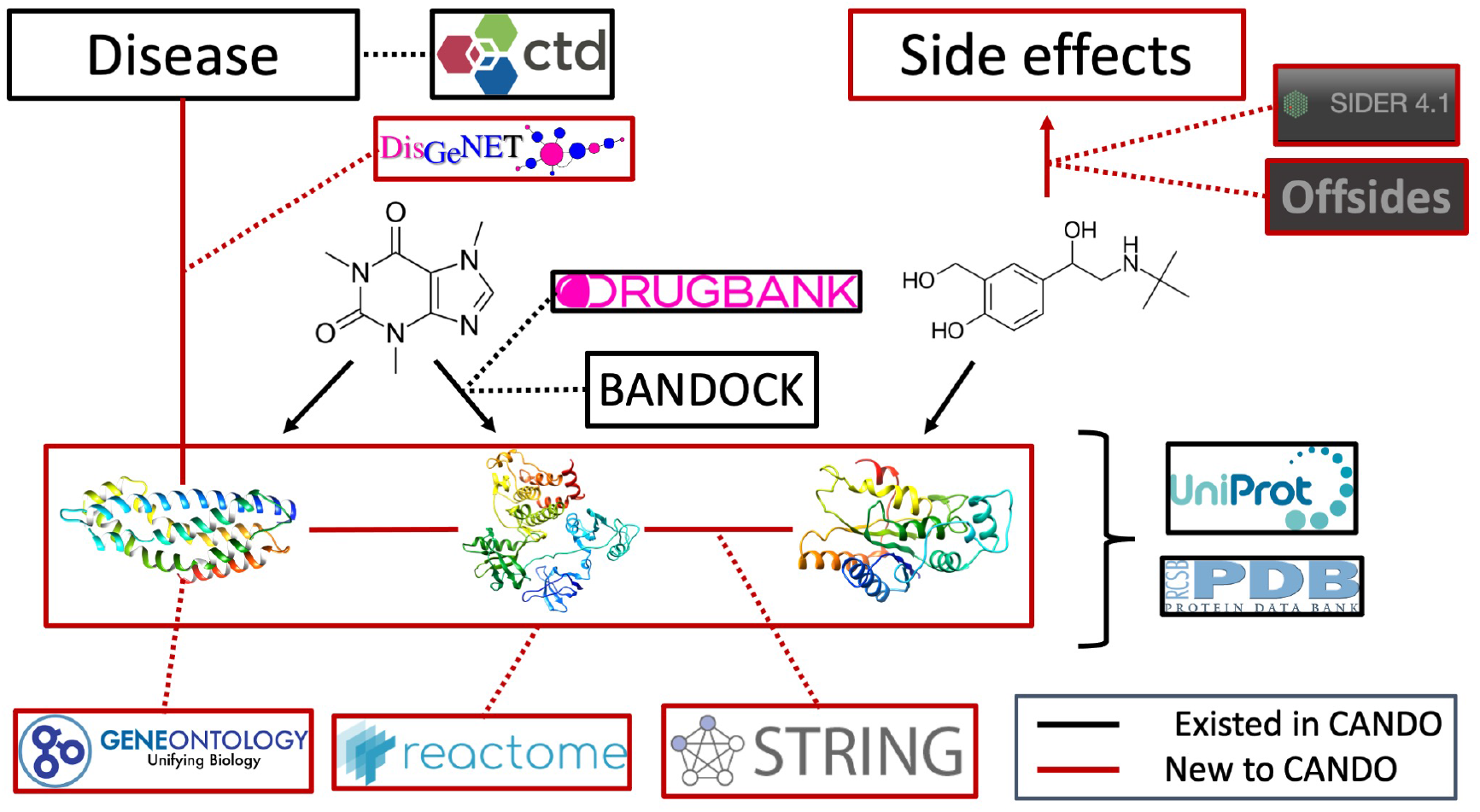
Overview of architecture and data sources for the heterogeneous network. Protein sequences and their x-ray structures were extracted from UniProt (Consortium, 2019b) and the Protein Data Bank (Burley et al., 2019), which were connected to each other via protein-protein interactions from the String database as well as to protein pathways from the Reactome database. Protein annotations to Gene Ontology (Consortium, 2004, 2015, 2019a) terms were extracted from UniProt as well. Proteins were mapped to indications using DisGeNET (Piñero et al., 2016). Drug/compound structures were downloaded from DrugBank (Wishart et al., 2017) in addition to known protein-drug interactions, which was further supplemented using our BANDOCK interaction scoring protocol (Falls et al., 2019; Mangione et al., 2020a). Finally, adverse drug reactions, or drug side effects, were extracted from the SIDER (Kuhn et al., 2016) and OFFSIDES (Tatonetti et al., 2012) databases. Black signifies an already existing entity or relation in the CANDO platform, whereas red signifies a novel association for our current network-based study and constitutes the majority of connections between these entities.

#### 4.1.1 Drug/compound library curation

13,194 drug and drug-like small molecules were extracted from DrugBank (Wishart et al., 2017) comprising 2,449 approved drugs, 2,519 metabolites, and the remaining 8,226 were either experimental, investigational, illicit, withdrawn, or veterinarian approved only compounds. Biologic therapeutics were excluded from the library. 337 compounds were removed following the protein-compound interaction scoring protocol (section 4.2) due to the lack of variance in their scores caused by their “simple” chemical composition, and were deemed uninteresting therapeutic candidates (such as elemental ions and inorganic salts). The final libraries used for this study consisted of either a 12,951 compound library following the addition of 24 compounds of interest implicated in indications for which CANDO has been applied (e.g. four GTPase KRAS inhibitors for non-small cell lung carcinoma), or the 2,336 approved drug sublibrary.

#### 4.1.2 Protein structure library curation

Protein structures for the human proteome were extracted from the Protein Data Bank (PDB) (Burley et al., 2019), if available, or modeled from sequence using I-TASSER version 5.1 (Zhang, 2008; Xu et al., 2011; Yang et al., 2015). Among the 19,582 human protein sequences extracted from UniProt (Consortium, 2019b), solved structures were chosen by (1) identifying all PDB entries that mapped to the UniProt identifier of interest using SIFTs (Dana et al., 2019; Velankar et al., 2012); (2) clustering them by the indices of their starting and ending sequence coordinates (relative to the full sequence) using the DBSCAN algorithm (Schubert et al., 2017); and (3) choosing the lowest resolution structures from each cluster that covered the largest portion of the full sequence while minimizing overlap (<50%). Further, solved chains were excluded if they were fewer than 100 residues. However, if the full sequence was fewer than 150 residues, this threshold was reduced to 30. This resulted in 4,966 proteins with at least one available solved structure in the PDB, comprising 5,316 total chains.

The remaining sequences were aligned to a library of PDB sequences with no greater than 70% sequence identity using the HHBLITS suite (Remmert et al., 2012). Structures were modeled in cases where sequence identity was ≤ 30 and the number of aligned residues was ≥ 100 if the sequence length was ≥ 150 residue else the number of aligned residues was ≥ 30. HHBLITS was run with one iteration. This resulted in 3,069 modeled structures for a total of 8,385 proteins comprising the human proteome.

#### 4.1.3 Indication/disease library curation

The CTD was used to map 22,772 drug-indication associations (Davis et al., 2021). All drug-indications associations were therapeutic in nature, i.e. the drug treats the underlying disease or condition. The indication library was further supplemented using the DisGeNET database (Piñero et al., 2016), which extracts protein-indication associations from existing databases as well as the literature using advanced text-mining methods. This added 4,961 indications and culminated in 337,177 protein-indication associations, which were split into four subsets according to their evidence-based scoring system thresholds of 0.0, 0.02, 0.1, 0.3, and 0.6. The evidence score is a culmination of the presence of the association in databases or the literature, as well as the consensus among these sources, with higher scores indicating greater confidence. Both databases utilize MeSH (Lipscomb, 2000) terms. The total count after combining the two data sources was 7,217 indications.

#### 4.1.4 Adverse drug reaction (ADR) library curation

ADRs, or drug side effects, were extracted from the SIDER database (Kuhn et al., 2016) which obtains them from drug labels; and the OFFSIDES project (Tatonetti et al., 2012), which used the US FDA’s Adverse Event Reporting System and Canada’s MedEffect resource to identify commonly reported ADRs for drugs that are not explicitly listed on their labels. Additionally, OFFSIDES used statistical techniques to identify ADRs for a drug that are not simply signs or symptoms of a disease for which a given drug is commonly indicated.

Of the 1,430 drugs present in SIDER, 686 were mapped to the CANDO compound library for a total of 92,143 associations to 5,144 ADRs. Due to the source of these associations being directly from drug labels, these associations were treated as a gold standard and were not further filtered.

Two separate versions of the OFFSIDES database were mapped to the CANDO drug library, with the first version sourcing ADR reports up to 2009, and the second version extending to 2014. After synthesizing these sets by matching the MedDRA ADR identifiers (Brown et al., 1999) of the second release to the Unified Medical Language System (UMLS) ADR identifiers (Bodenreider, 2004) of the first release, 1,622 drugs in CANDO were associated to 10,268 unique ADRs for a total of 191,591 associations.

Further filtering of this set was necessary due to the significant number of drugs with contradictory ADR and disease associations. A trivial example of this would be if a given drug was associated to the ADR “nausea” from SIDER/OFFSIDES, implying the drug *induces* nausea, but was also associated to the equivalent term for “nausea” in the CTD, implying the drug *treats* nausea. To prevent biasing the network by including these drug-ADR associations that could artificially enhance drug-indication prediction performance, all ADRs and indications were first mapped to their equivalent SNOMED (Donnelly et al., 2006) clinical terminology concepts using the UMLS metathesaurus, and only ADR concepts belonging to the SNOMED “disorder” class were kept.

For a given drug, ADR associations were removed if the UMLS identifier of the ADR term mapped to the same SNOMED concept as any of the MeSH identifiers for the indications the drug is known to treat. Further, for a given indication and its associated drugs, all of the other indications associated to those drugs were grouped; if any of the mapped SNOMED concepts for these indications matched the mapped SNOMED concept for an ADR, associations to this ADR for all these drugs were removed.

This process was repeated for all indications in CANDO with the indication grouping scheme serving as a proxy for drug therapeutic class. Lastly, the concept hierarchy from SNOMED was used to perform a filtering process that utilized “is_a” relations to connect the lower level ADR concepts to their upper level classes (e.g. both “Low blood pressure” and “Heart disease” belong to “Disorder of cardiovascular system”). Similarly to the process used for indications, all mapped SNOMED concepts of the ADRs and indications that belonged to same upper level SNOMED concept were grouped; if a drug was associated to both an indication and ADR of the same upper level class (excluding top level terms “Disease” and “Clinical finding”), the ADR association was removed. This culminated in 50,210 associations between 1,112 drugs and 1,413 ADRs, after removing all ADRs with fewer than 10 drugs associated. This was combined with the 85,095 associations from SIDER that could be mapped to SNOMED for a final, non-redundant set of 133,322 associations between 4,245 ADRs and 1,171 compounds.

#### 4.1.5 Protein-protein and protein-pathway association curation

All human proteins were associated with other proteins in terms of their known interactions and the biological pathways to which they belong using the STRING and Reactome databases (Jassal et al., 2020; Szklarczyk et al., 2019) respectively. STRING collates protein-protein interactions from multiple sources, including the literature and other databases, evolutionary information transferred from known protein-protein interactions in other organisms, computational predictions, and other data sources (Szklarczyk et al., 2019). A cumulative score is assessed based on the available evidence for a protein pair; in this study, a cutoff of 700 was used to determine a protein-protein interaction, which is deemed high confidence according to the curators, resulting in 169,208 interactions. Reactome constituted 2,219 pathways comprising 108,466 associations, with an average of ≈49 structures per pathway.

#### 4.1.6 Gene Ontology annotation curation

Annotations to the Gene Ontology (GO) for all human proteins as provided by the UniProt database were extracted for all descendant terms of the three main upper level entities (“biological process”, “molecular function”, and “cellular component”). This resulted in 18,094 unique GO entities that gave rise to 262,059 total annotations.

### 4.2 Compound-protein interaction calculation

Interaction scores between each protein and compound structure were computed using our in-house bioanalytic docking protocol BANDOCK (Falls et al., 2019; Mangione et al., 2020a). BANDOCK uses a library of protein structures with known ligands bound from the PDB determined using x-ray diffraction to assign a real-value interaction score between a compound and protein for a target structure based on the similarity principle (i.e., via homology inference). A real value interaction score is assigned between a compound and a protein using the latter’s known or predicted binding sites derived from its structure, and molecular fingerprinting to determine compound-compound similarity based on presence or absence of chemical substructures. The binding site prediction component is handled by the COACH algorithm of the I-TASSER version 5.1 suite (Yang et al., 2013), which utilizes a consensus scoring approach of three different methods to score how similar the binding sites on the query protein are to the binding sites in our known protein-compound structure library. The output of COACH is a list of binding sites, the ligands known to bind to them, and an associated confidence score (Pscore) for each site. To assign a score for a compound-protein interaction, the Sorenson-Dice coefficient (Duarte et al., 1999) between the ECFP4 chemical fingerprints (Rogers and Hahn, 2010) of each ligand associated to the predicted binding sites for the protein and the query compound (Cscore) are computed; the strongest matching ligand (Cscore closest to 1.0) is chosen and multiplied by the Pscore for the binding site to which this ligand is associated (Pscore × Cscore). This process is iterated for all compound and protein pairs to generate a proteomic interaction signature for every drug/compound in the corresponding library. The interaction signature is simply a real value vector interaction scores that we hypothesize to describe compound behavior.

### 4.3 All-against-all (shotgun) drug repurposing benchmarking

The main goal of the CANDO platform is to accurately describe drug/compound behavior, and the primary protocol used to assess the performance of a particular pipeline utilizes the indication approvals extracted from the CTD to determine how often drugs associated to the same indication are predicted accurately (i.e., most similar) according to various thresholds/cutoffs. In the traditional pipeline, the similarity of two compounds is calculated by comparing the scores in their proteomic interaction signatures using the root mean square deviation (RMSD) metric. This is repeated iteratively for all pairs in the library, resulting in each having a list of compounds ranked by decreasing similarity. For each drug approved for a given indication, the benchmarking protocol averages how many times another drug associated to that indication is found within a certain rank cutoff (10, 25, 100, etc) in the list of most similar compounds to the hold-out drug; a topN accuracy is assessed for the indication based on the number of times another associated drug was found in the top *N* most similar compounds for each drug. This is repeated for all indications to determine the global topN accuracy, and then repeated for all other cutoffs. This metric is known as the average indication accuracy; the average pairwise accuracy and the indication coverage are also computed which are the weighted average based on how many drugs are associated with each indication and the number of indications with non-zero accuracies, respectively (Minie et al., 2014; Mangione et al., 2020a; Schuler et al., 2021). For this study, the benchmarking protocol only included the sublibrary of 2,336 approved drugs and used cosine distance as opposed to RMSD as it provided a substantial increase in computational speed for the similarity calculations with no change in performance.

### 4.4 Compound-protein interaction representation within the networks

Deciding which of the > 100 million interaction scores between every protein and compound in CANDO to include in our networks is a non-trivial problem. This is in part due to the unweighted, binary nature of the edges in the network architecture. Further, smaller compounds tend to be favored by the ECFP4 fingerprint, while many others structurally resemble ligands commonly co-crystallized with proteins in the PDB, both of which inflate their similarity scores (Cscores). In addition, well-studied proteins like cancer targets or kinase families also tend to be overrepresented in the PDB and therefore have greater binding site similarity scores (Pscores) from COACH, as well as increased ligand partner diversity as a consequence. Since the Pscore from COACH implicitly measures confidence in the binding site and therefore its associated ligand, simply considering only the Cscore (as opposed to multiplying it by the Pscore) during the BANDOCK scoring protocol would disregard a significant portion of the embedded evolutionary information and reduce the discernability between a compound with the same matching ligand for multiple proteins (section 4.2). Therefore, a normalization scheme was devised to select the strongest and most unique interactions between all compounds and proteins.

For a given interaction score between a compound and protein, the min-max normalized score was computed based on the non-zero minimum and maximum values among all interaction scores for the corresponding protein; this process was iterated for all proteins in the compound signature. The normalized scores were then ranked for the compound and the top five interactions were initially selected to be included in the network. If a tie occurred at ranks five and six, all subsequent interactions equal to the value at rank five were also included. This process was repeated for every compound in the library. To further reduce any bias that may still be present after normalization, all proteins were reduced to their top 50 compounds; if ties occurred at rank 50, all tying compounds were ranked by the inverse of their average interaction score against all proteins, which therefore prioritized more unique interactions. This culminated in 50,345 interactions across 6,161 (73.5%) proteins.

Compound-protein interactions annotated in DrugBank were also included in the network, constituting 13,702 known interactions, of which 1,876 (13.7%) were captured in the normalization scheme. This resulted in 62,510 unique compound-protein associations after combining the interactions from multiple chains of the same protein.

### 4.5 Network integration and preliminary analyses

The following biological entities and relations were initially included in the network for preliminary analysis: drugs/compounds, proteins, indications, pathways, GO terms, ADRs, protein-protein interactions, compound-protein interactions, compound-ADR associations, protein-pathway associations, and protein-GO annotations (Figure 6). Known drug-indication approvals were excluded from all networks prior to benchmarking. The complete network featured 64,308 nodes and 1,072,280 edges and was built using the Python package NetworkX (Hagberg and Conway, 2020). Separate networks were built which either included or excluded each of the relation types above, except for the protein-indication associations in which the five DisGeNET thresholds were varied instead. This resulted in a total of 80 unique networks.

For each network architecture, node representations/embeddings were generated using the Python implementation of the node2vec algorithm (Grover and Leskovec, 2016), which uses biased random walks starting from each node to produce a continuous value vector in a lower dimensional space that aims to preserve the contextual neighborhood in which the node is situated. The walk length, number of walks, and output dimensionality were set to 10, 100, and 128, respectively, for all networks. To assess how well the node embeddings captured the therapeutic potential of compounds in the network, the embedded vectors (aka multiscale or interactomic signatures) for the approved drugs were used as input to the benchmarking protocol in place of their traditional protein interaction signatures. Drug-drug similarities and drug repurposing accuracies were computed as described above (section 4.3). For this analysis, a drug-drug similarity matrix was also computed using the ECFP4 chemical fingerprint to use as a control comparison.

### 4.6 Adverse drug reaction prediction

ADRs were predicted for all 12,951 compounds in the CANDO library based on the similarity of each compound to drugs with known ADRs. Signature similarity is first computed for all compound-compound pairs in the library in the same way as for the benchmarking protocol (section 4.3). The ADRs associated with the most similar compounds to a query compound within a set cutoff *N* are counted; a probability is assessed based on the number of times the ADR was present, the frequency of the ADR within the library, the size of the compound library, and the cutoff N, using a hypergeometric distribution. The ADRs are ranked by the probability, which represents the chance of randomly selecting at least that many compounds associated with the ADR without replacement in *N* trials.

### 4.7 Pathway-disease association extraction

The importance of each pathway in predicting drug-indication associations was assessed via random forest machine learning models. For 60 indications with at least 50 associated drugs based on the CTD mapping, a random forest model was trained using 90% of the associated drugs as the positive samples, the remaining 10% as the positive test samples, and an equal number of training and test “negative” samples chosen at random from the sublibrary of drugs with at least one indication association, ensuring that the randomly chosen drug is not already associated to the indication or any upper level class to which the indication belongs. The features of the drugs were a binary vector of integers with true (1) representing the distance of the drug to the pathway in the network being less than or equal to three, otherwise false (0). This ensured that the drug directly interacted with a protein in the pathway (distance of two), or interacted with an ancillary protein that shared an interaction with a protein in the pathway (distance of three). The pathways were filtered for those that satisfied the following criteria: (1) at least 5 and no greater than 250 proteins associated; 2) at least two associated proteins with at least one interaction to any compound in the network; and 3) ≥ 1% and ≤ 33% of compounds with a distance of two to the pathway. Ten iterations were performed for each indication; the average performance over all iterations was assessed and the feature importances for the indication models with the best average performance were extracted.

## 5 CONCLUSION

This study is the first using the CANDO platform for large-scale integration of higher level biomedical processes and entities such as protein pathways, protein-protein interactions, ADRs, and protein-indication associations, with the primary goal of better understanding the totality of interactions, or the multiscale interactomic signatures, of drugs in relation to not only indications that they are approved for, but also the ADRs that they are associated with. The rich signal provided by the ADRs for describing drug behavior far outweighed the contributions of other relations in terms of drug-indication benchmarking, which will be further explored in future studies. However, the interactomic approach was still able to accurately predict drug-indication associations when drugs were considered by only their distance to a subset of the pathways in the network, indicating the contextual information of the protein interactions is still relevant for describing their behavior. Further, the networks constructed without any ADRs still outperformed the linear proteomic signature pipeline, albeit to a much lesser extent, providing further justification for our multiscale interactomic approach to advance the science of drug discovery.

## AUTHOR CONTRIBUTIONS

W.M. conceived the prediction pipelines, research design, approach and methods, conducted all experiments and analysis, implemented all pipelines, and drafted the manuscript. Z.F. helped with data generation, research design, approach, and methods, and editing the manuscript. R.S. conceived the prediction pipelines, research design, approach and methods, supervised the overall project, and edited the manuscript. All authors have read and agreed to the published version of the manuscript.

## ACKNOWLEDGMENTS

The authors would like to acknowledge the support provided by the Center for Computational Research at the University at Buffalo. We would also like to thank all members of the Samudrala Computational Biology Group.

## DATA AVAILABILITY STATEMENT

The datasets generated for this study can be found on the Samudrala Group website http://compbio.buffalo.edu/data/mc_cando_interactomics/.

